# 65°C Short-Term Heating: A Method for Transporting Urine Samples Over Long Distances Without Cold Chain

**DOI:** 10.64898/2025.12.31.697171

**Authors:** Yuzhen Chen, Youhe Gao

## Abstract

Biomarkers are measurable changes associated with physiological or pathophysiological processes in the body. Urine, not strictly regulated by homeostatic mechanisms, accumulates extensive variable information and reflects changes in the body earlier and more sensitively, making it an excellent source for biomarker discovery. However, long-distance transport of urine samples usually requires a strictly maintained cold chain throughout the process, resulting in high costs and operational challenges. Here, we propose a urine sample processing method that uses brief heating to inhibit bacterial proliferation and protein degradation. A single urine sample was equally divided into three groups: a pre-heated transport group, a non-heated transport group, and a -80°C frozen storage control group, with five replicates in each group. Both the pre-heated and non-heated transport groups were transported simultaneously across northern and southern China for five days under ambient temperature conditions. Samples were analyzed using liquid chromatography coupled with tandem mass spectrometry. The results showed that the intra-group overlap rates of identified proteins within the groups treated with 65°C water bath heating for 15 minutes and -80°C frozen storage were 97.1% and 97.5%, respectively, demonstrating high reproducibility. There were no significant differences in the number or types of urinary proteins identified between these two groups, with an inter-group overlap rate of 99.9%. Additionally, the pre-heated transport group and frozen storage control group showed no significant difference in the number of bacterial-derived peptides and proteins identified. In contrast, significantly more bacterial-derived peptides and proteins were identified in the non-heated transport group than in both the pre-heated transport and frozen storage control groups. This indicates that heating effectively suppresses bacterial counts in urine samples during five days of transport at ambient temperatures. This approach provides a more economical and convenient solution for long-distance transport of urine samples, eliminating the need for cold chain transportation or preservatives. We also recommend using consistent urine processing methods within the same study to minimize technical variability.

## 1 Introduction

Biomarkers are measurable changes associated with physiological or pathophysiological processes in the body, playing an important role in clinical diagnosis, treatment and prognosis. Biological samples obtained from patients constitute important resources for medical research and clinical translation. Among these samples, urine, as a filtrate of blood, does not need or possess homeostatic mechanisms to be stable. It accommodates and accumulates more changes without harming the body, reflecting changes in all organs and systems of the body earlier and more sensitively. Therefore, urine represents an excellent source of biomarkers [1].

Ensuring “high-quality” biological samples is critical [2]. Previous studies indicated that, to prevent protein degradation, urine should be stored at low temperatures shortly after collection and avoid repeated freeze-thaw cycles [3-5]. In some studies, protease inhibitors are added during urine collection, particularly in cases of proteinuria, but these inhibitors themselves interfere with subsequent proteomic analysis [5, 6]. In addition, bacterial overgrowth significantly alters the urine proteome, posing a major challenge for sample collection and storage. To prevent bacterial overgrowth, preservatives such as sodium azide or boric acid are recommended in many proteomic analysis protocols [7, 8], especially when urine is stored at room temperature for more than 8 hours or at 4°C for longer than 16 hours [5].

Our laboratory previously proposed a method in which urinary proteins are adsorbed onto polyvinylidene fluoride (PVDF) or nitrocellulose (NC) membranes and then dried for storage. This approach effectively conserves storage space and enables the preservation of large-volume, high-volume urinary proteins [9]. However, this method is limited by the need for specialized equipment and reagents, and typically requires trained personnel for operation. When this membrane-based preservation method is not adopted, long-distance transport of urine samples requires strict maintenance of cold chain to keep the samples frozen throughout the process, which is costly and operationally inconvenient. Inadequate freezing can easily lead to protein degradation and bacterial overgrowth, thereby compromising experimental results. Herein, we propose a novel urine sample processing method that employs heat treatment to inhibit bacterial proliferation and protein degradation, with the aim of enabling a more economical and convenient approach for the long-distance transport of urine samples (Figure 1).

**Figure 1.**
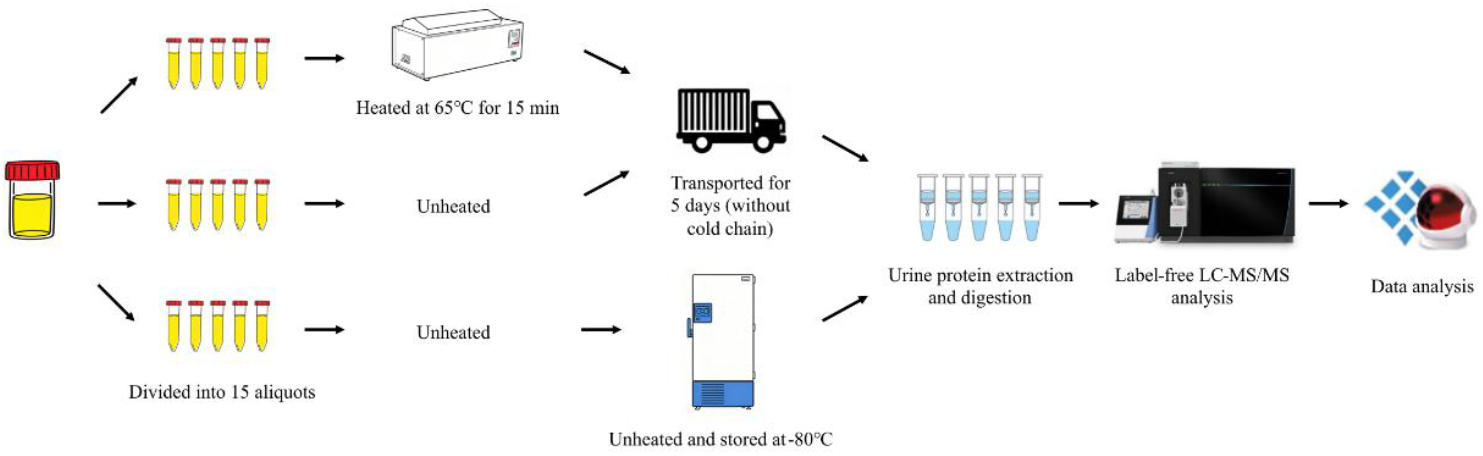
Technical route for exploring methods of long-distance transport of urine samples without cold chain.

## 2 Materials and Methods

### 2.1 Urine Sample Pretreatment

Approximately 200 mL of urine samples were collected from a healthy volunteer, and centrifuged at 12 000 ×*g* for 20 min at 4°C. The supernatant was evenly divided into 15 aliquots. Five samples were directly stored at -80°C as the frozen storage control group. Another five samples were heated in a 65°C water bath for 15 min, and then transported together with the remaining five unheated samples via SF Express Standard Service. The transport route spanned northern and southern China. Samples were dispatched from Beijing on July 15, 2025 (ambient temperature range: 23-31°C), transported to Shantou, Guangdong (ambient temperature range on arrival: 28-33°C), and then returned to Beijing under the same conditions, arriving on July 20 (ambient temperature range: 23-29°C). The entire transportation process lasted 5 days. No cold chain or special temperature control measures were used during transportation. All 10 samples were exposed to ambient temperature conditions throughout the transport period.

### 2.2 Urine Sample Preparation for Label-free Analysis

Upon receiving the samples, 20 mmol/L dithiothreitol solution (DTT, Sigma) was added, vortexed, and mixed well. The samples were heated in a metal bath at 37°C for 1 h. After cooling to room temperature, 50 mmol/L iodoacetamide solution (IAA, Sigma) was added, vortexed, and mixed well, and reacted at room temperature without light for 40 min. Pre-cooled anhydrous ethanol at four times the volume of the sample was added, mixed thoroughly, and then precipitated at -20°C for 36 h. Following centrifugation at 10 000 ×*g* for 30 min at 4°C, the supernatant was discarded. The protein precipitate was then suspended in an appropriate lysis buffer (8 mol/L urea, 2 mol/L thiourea, 25 mmol/L dithiothreitol, and 50 mmol/L Tris) to obtain urine protein extract. Protein concentration was quantified using the Bradford kit assay (Applygen, Beijing, China).

Using filter-aided sample preparation (FASP) method [10], a total of 100 μg of protein was added to a 1.5 mL centrifuge tube. UA solution (8 mol/L urea, 0.1 mol/L Tris-HCl, pH 8.5) was added to make a total volume of 200 μL. 200 μL of UA solution was added to a 10 kD ultrafiltration tube (Pall, Port Washington, NY, USA) and centrifuged twice at 14 000 ×*g* for 10 min at 18°C. The treated protein sample was added and centrifuged at 14 000 ×*g* for 40 min at 18°C. 200 μL of UA solution was then added and centrifuged at 14 000 ×*g* for 40 min at 18°C, repeated once. 25 mmol/L NH_4_HCO_3_ solution was added and centrifuged at 14 000 ×*g* for 40 min at 18°C, repeated once. The samples were digested overnight at 37°C with trypsin (Trypsin Gold, Promega, Madison, WI, USA) at an enzyme-to-protein ratio of 1:50. The digested peptides were eluted from ultrafiltration membranes, desalted with HLB columns (Waters, Milford, MA), dried in a vacuum desiccator, and stored at -80°C.

### 2.3 Liquid Chromatography Coupled with Tandem Mass Spectrometry Analysis

The digested peptides were dissolved in 0.1% formic acid, and the peptide concentration was quantified using a BCA kit. The peptides were then diluted to a final concentration of 0.5 μg/μL. The indexed retention time (iRT) reagent (Biognosis, Schlieren, Switzerland) was then added at a ratio of 1:10 (iRT-to-sample). 1 μg of the peptide from each sample was separated using the EASY-nLC 1200 system (Thermo Fisher Scientific, Waltham, MA, USA) and analyzed with an Orbitrap Fusion Lumos Tribrid mass spectrometer (Thermo Fisher Scientific, Waltham, MA, USA). Individual samples were analyzed in data-independent acquisition (DIA) mode, with each sample run in triplicate. After every 8-9 runs, a single DIA analysis of the pooled peptides was performed to control the quality of the whole analytical process.

The sample was loaded onto a reversed-phase C18 trap column (75 μm × 2 cm, 3 μm) at a flow rate of 0.4 μL/min and separated with a reversed-phase analytical column (75 μm×25 cm, 2 μm). A 90-minute gradient elution was applied using mobile phase A (0.1% formic acid) and mobile phase B (0.1% formic acid in 80% acetonitrile). The spray voltage was set to 2.5 kV. A full MS scan was acquired within a 400-1 200 *m/z* range with a resolution of 120 000. The MS/MS scan was acquired in Orbitrap mode in the range of 200-2 000 *m/z* with a resolution of 30 000. The HCD collision energy was set to 32%.

### 2.4 Database Searching and Data Processing

The results were then imported into Spectronaut Pulsar (version 19, Biognosys AG, Schlieren, Switzerland) software for analysis and processing. The peptide intensity was calculated by summing the peak areas of the respective fragment ions for MS^2^, while protein intensity was calculated by summing the peptide intensities. Total proteins were identified based on the criteria that each protein contained at least two specific peptides and a false discovery rate (FDR) <1% at the protein level. Protein identification results were compared among the three groups: the pre-heated transport group, the non-heated transport group, and the -80°C frozen storage control group.

## 3 Results and Discussion

### 3.1 Heat Treatment at Different Temperatures

To determine the optimal experimental conditions, we first designed a bivariate protocol consisting of 7 temperature gradients, with two storage durations assigned to each gradient, together with a frozen storage control group. A single urine sample was collected, centrifuged, and the supernatant was divided into 15 aliquots. Among these, one aliquot was stored directly at -80°C, while the remaining 14 aliquots were subjected to different heating treatments: water bath heating at 60°C for 30 min, 65°C for 15 min, 70°C for 15 min, 75°C for 10 min, 80°C for 10 min, and 90°C for 5 min, as well as metal bath heating at 99.9°C for 5 min, with two aliquots assigned to each temperature condition. For each temperature condition, one aliquot was stored at ambient temperature for 3 days and the other for 5 days. LC-MS/MS analysis showed that the numbers of identified peptides and proteins in all treatment groups did not differ significantly from those in the frozen storage control group (Table 1).

**Table 1.** Number of identified peptides and proteins in individual samples.

| Sample No. | Treatment | Ambient temperature<br>storage duration/d | Number of<br>identified peptides | Number of<br>identified proteins |
| --- | --- | --- | --- | --- |
| 1 | Heated at 60°C for 30 min | 3 | 24484 | 2892 |
| 2 | Heated at 60°C for 30 min | 5 | 24448 | 2868 |
| 3 | Heated at 65°C for 15 min | 3 | 25227 | 2922 |
| 4 | Heated at 65°C for 15 min | 5 | 25185 | 2907 |
| 5 | Heated at 70°C for 15 min | 3 | 24861 | 2914 |
| 6 | Heated at 70°C for 15 min | 5 | 19109 | 2804 |
| 7 | Heated at 75°C for 10 min | 3 | 19354 | 2820 |
| 8 | Heated at 75°C for 10 min | 5 | 19921 | 2824 |
| 9 | Heated at 80°C for 10 min | 3 | 25712 | 2942 |
| 10 | Heated at 80°C for 10 min | 5 | 25645 | 2925 |
| 11 | Heated at 90°C for 5 min | 3 | 25598 | 2937 |
| 12 | Heated at 90°C for 5 min | 5 | 25765 | 2942 |
| 13 | Heated at 99.9°C for 5 min | 3 | 25831 | 2950 |
| 14 | Heated at 99.9°C for 5 min | 5 | 25897 | 2945 |
| 15 | Unheated | Stored at -80°C | 24576 | 2941 |

### 3.2 Identification of Urinary Proteins

Based on the preliminary results, water bath heating at 65°C for 15 min was selected for the formal experiment. A single urine sample was equally divided into three groups: a pre-heated transport group, a non-heated transport group, and a -80°C frozen storage control group. Both the pre-heated and non-heated transport groups were transported under identical conditions, with samples exposed to ambient temperatures throughout the process. In total, 15 samples from the three groups were analyzed by liquid chromatography coupled with tandem mass spectrometry. The results showed that the numbers of identified peptides and proteins in the pre-heated transport group were not obvious different from those in the frozen storage control group. In contrast, the non-heated transport group exhibited lower numbers of identified peptides and proteins than the frozen storage control group (Table 2).

**Table 2.**
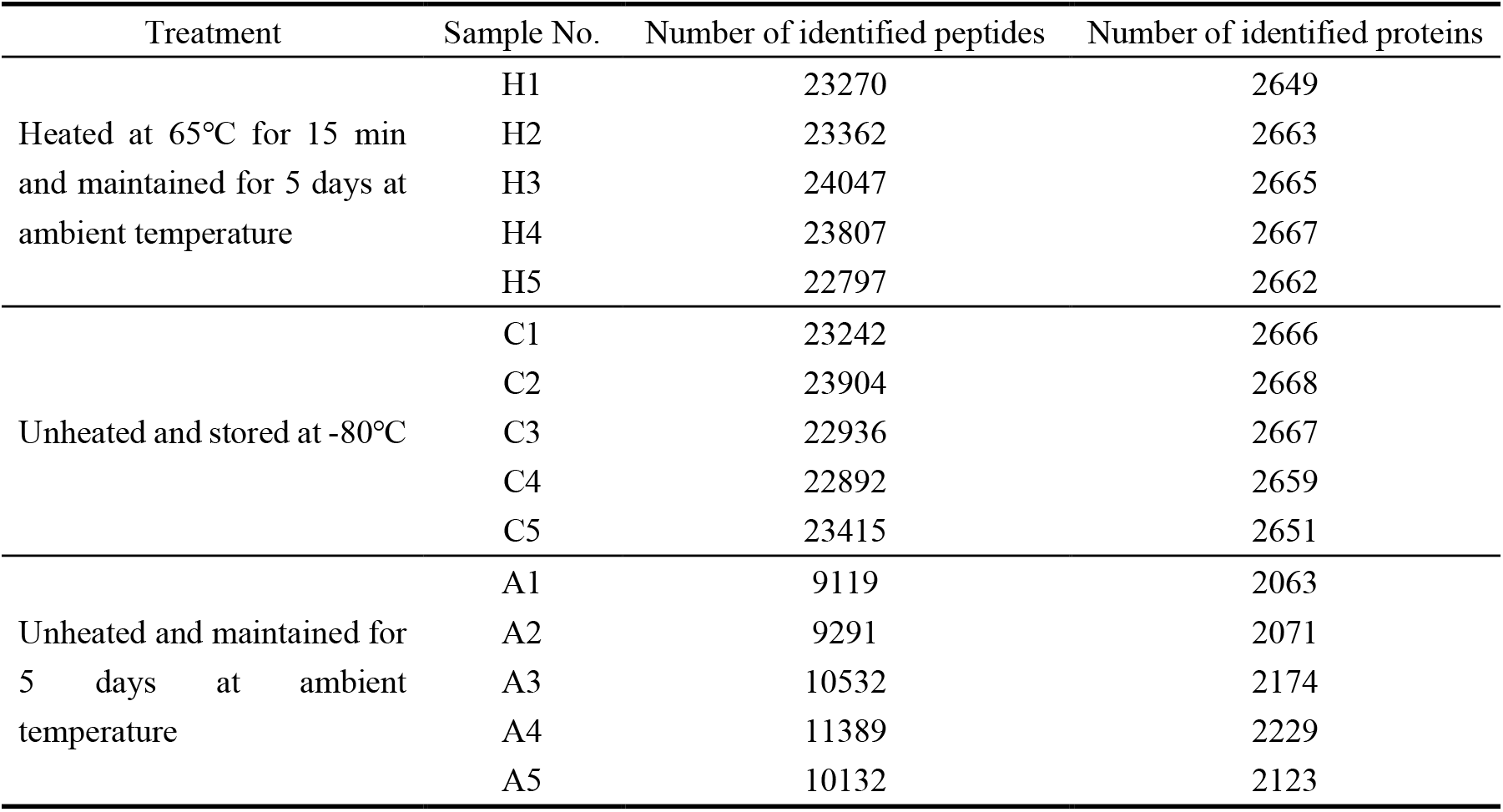
Number of identified peptides and proteins in individual samples (search results from the *Homo sapiens* database).

Principal component analysis (PCA) was performed on the total proteins using the SRplot web server (http://www.bioinformatics.com.cn/) (Figure 2). The results showed no significant difference between the pre-heated transport group and the frozen storage control group, whereas the non-heated transport group was clearly distinguishable from both the pre-heated transport group and the frozen storage control group.

**Figure 2.**
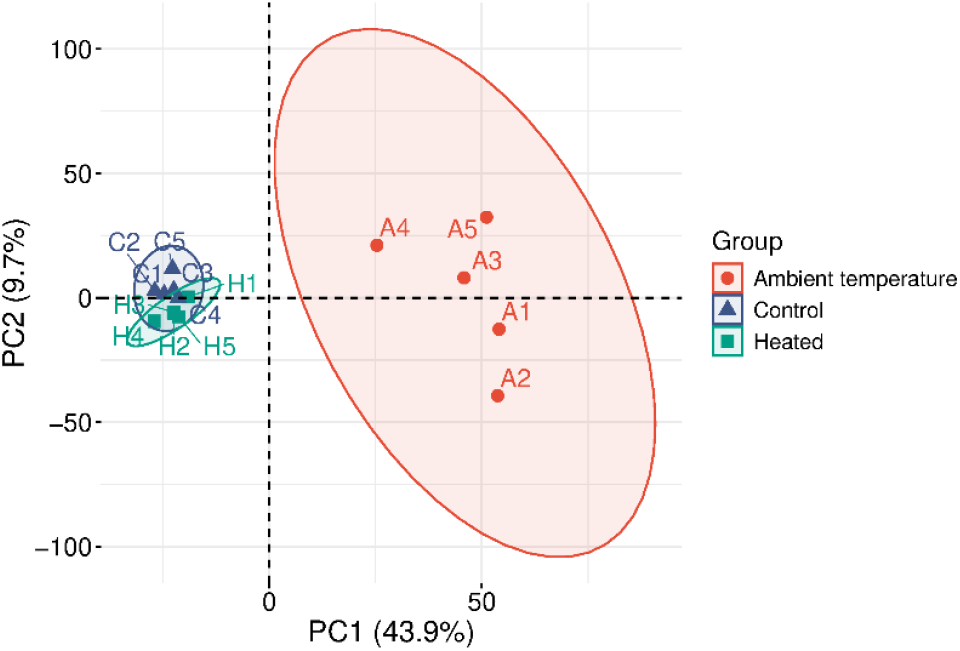
Principal component analysis of the total proteins.

### 3.3 Protein Overlap Rate Identified by Mass Spectrometry

#### 3.3.1 Intra-Group Overlap Rate

Protein identification results and intra-group overlap rates were calculated for the pre-heated transport group, the frozen storage control group, and the non-heated transport group (intra-group overlap rate = number of shared proteins / total number of identified proteins ×100%) (Table 3). Additionally, Venn diagrams were used to visualize the intra-group overlap of identified proteins within each group (Figure 3). The results showed that the intra-group overlap rates of the pre-heated transport group and the frozen storage control group were 97.1% and 97.5%, respectively, indicating high intra-group repeatability in both groups. In contrast, the total number of identified proteins in the non-heated transport group was lower than that in the pre-heated transport group and the frozen storage control group, and its intra-group overlap rate was only 74.2%. This markedly lower overlap rate suggests poor intra-group reproducibility in the non-heated transport group.

**Table 3.**
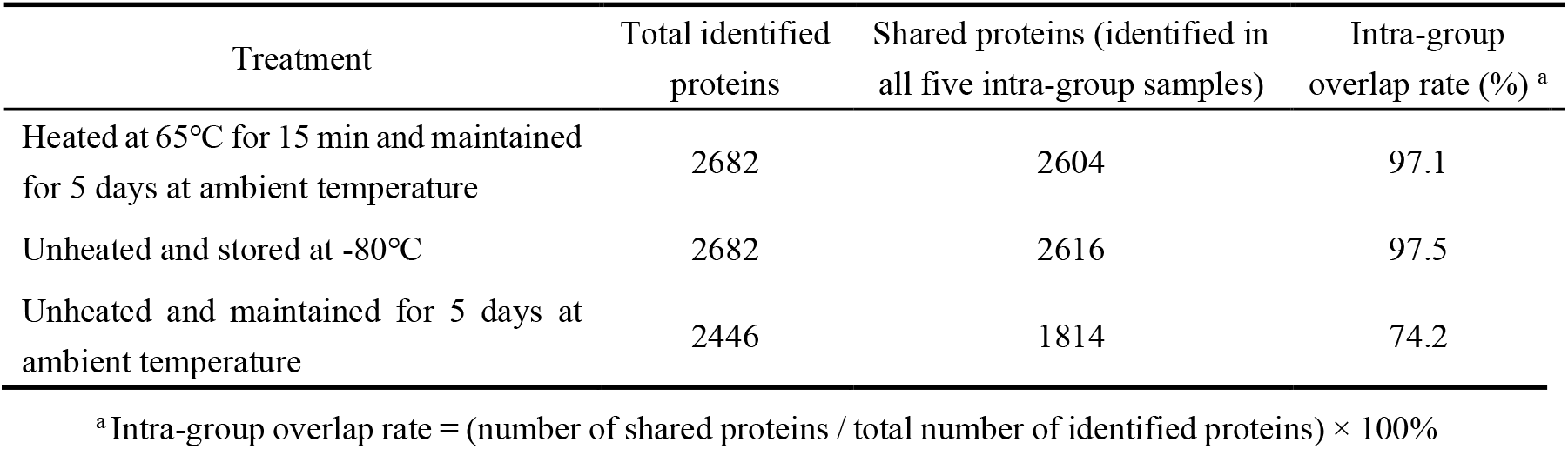
Statistics of protein identification and overlap rates in each group.

**Figure 3.**
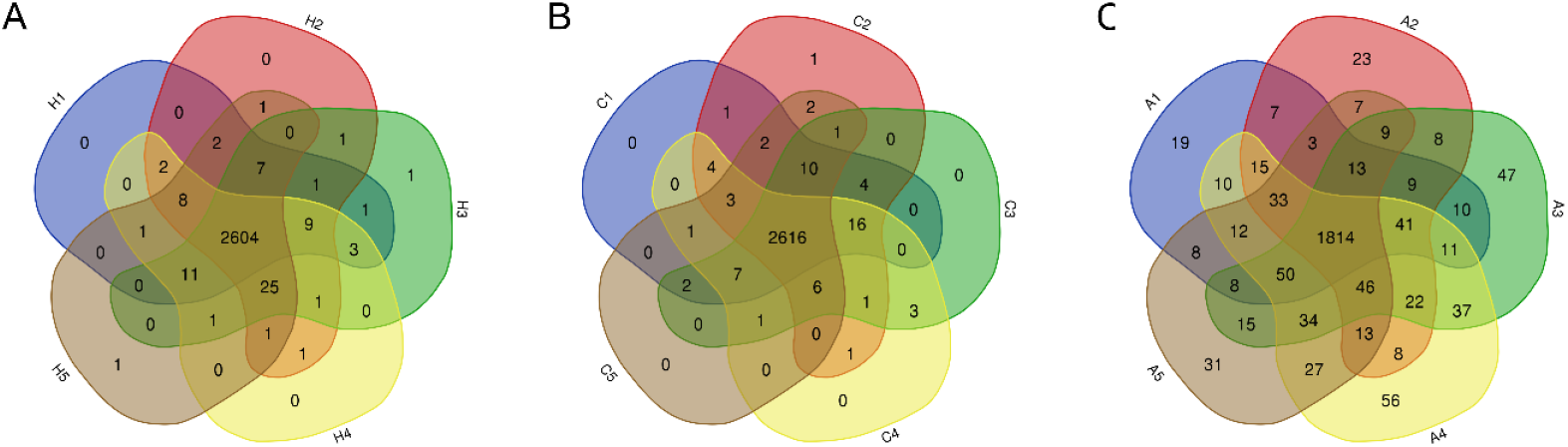
Intra-group overlap of identified proteins. (**A)** Pre-heated transport group; (**B)** Frozen storage control group; (**C)** Non-heated transport group.

#### 3.3.2 Inter-group Overlap Rate

A Venn diagram was also used to visualize the inter-group overlap of identified proteins among the three groups (Figure 4). A total of 2 680 proteins were commonly identified in the pre-heated transport group and the frozen storage control group, corresponding to an inter-group overlap rate of 99.9% (inter-group overlap rate = number of shared proteins / total number of identified proteins ×100%). In contrast, the non-heated transport group exhibited an overlap rate of 90.8% with both the pre-heated transport group and the frozen storage control group, respectively, which was lower than that observed between the other two groups.

**Figure 4.**
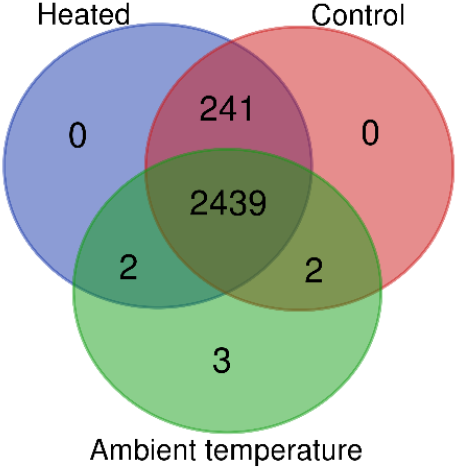
Inter-group overlap of identified proteins.

### 3.4 Bacterial Database Search Results

Samples from the non-heated transport group appeared visibly turbid, whereas samples from the pre-heated transport group transported under the same conditions showed no turbidity, consistent with the appearance of the frozen storage control group. Previous studies have suggested that preservatives such as sodium azide and boric acid should be added to urine stored at room temperature for more than 8 hours to prevent bacterial overgrowth [5]. To investigate whether heat treatment could effectively control bacterial counts in urine, the data acquired using the DIA mode were imported into Spectronaut Pulsar software, and searched against a bacterial database downloaded from the UniProt database (updated in August 2025). The results showed that the numbers of bacteria-derived peptides and proteins identified in the pre-heated transport group did not differ significantly from those in the frozen storage control group. In contrast, the non-heated transport group exhibited significantly higher numbers of bacteria-derived peptides and proteins than the frozen storage control group (Table 4). These findings indicate that heating urine samples in a 65°C water bath for 15 min can effectively control the bacterial count during 5 days of transport at ambient temperature.

**Table 4.** Number of identified peptides and proteins in individual samples (search results from the bacterial database).

| Treatment | Sample No. | Number of identified peptides | Number of identified proteins |
| --- | --- | --- | --- |
| Heated at 65°C for 15 min and maintained for 5 days at ambient temperature | H1 | 40 | 6 |
|  | H2 | 30 | 9 |
|  | H3 | 59 | 12 |
|  | H4 | 48 | 16 |
|  | H5 | 47 | 13 |
| Unheated and stored at -80°C | C1 | 44 | 24 |
|  | C2 | 31 | 7 |
|  | C3 | 51 | 11 |
|  | C4 | 74 | 18 |
|  | C5 | 67 | 12 |
| Unheated and maintained for 5 days at ambient temperature | A1 | 14138 | 1834 |
|  | A2 | 14277 | 1857 |
|  | A3 | 14353 | 1839 |
|  | A4 | 14344 | 1833 |
|  | A5 | 14434 | 1856 |

Previous studies on the thermal resistance of common water-borne or food-borne pathogenic and indicator bacteria at 55-65°C (a temperature range relevant to domestic hot water systems) have shown that this range is critical for effectively eliminating intestinal pathogenic bacteria and their components. At 65°C, the *D*-value (the time required to reduce the bacterial population by 90%) of *Enterococcus faecalis* is 7-19 s, whereas the *D*-values of other tested strains, including *Escherichia coli, Pseudomonas aeruginosa*, and *Serratia marcescens*, are all less than 6 s [11]. This further supports the effectiveness of heat treatment as a method for controlling bacterial counts.

## 4 Conclusion

The urine sample processing method involving water bath heating at 65°C for 15 min demonstrates excellent reproducibility, and shows no significant differences from the conventional -80°C frozen storage method in terms of the number and types of identified urinary proteins. This approach provides a more economical and convenient strategy for the long-distance transport of urine samples. In addition, we recommend that a consistent urine processing method be used within the same study to minimize technical variability.

